# A detailed spatio-temporal atlas of the white matter tracts for the fetal brain

**DOI:** 10.1101/2024.04.26.590815

**Authors:** Camilo Calixto, Matheus Dorigatti Soldatelli, Camilo Jaimes, Simon K. Warfield, Ali Gholipour, Davood Karimi

**Affiliations:** Computational Radiology Laboratory (CRL), Boston Children’s Hospital, Harvard Medical School; Division of Neuroradiology, Department of Radiology, Boston Children’s Hospital, Harvard Medical School; Massachusetts General Hospital, 55 Fruit St, Boston, MA 02114, USA April 25, 2024

## Abstract

This study presents the construction of a comprehensive spatiotemporal atlas detailing the development of white matter tracts in the fetal brain using diffusion magnetic resonance imaging (dMRI). Our research leverages data collected from fetal MRI scans conducted between 22 and 37 weeks of gestation, capturing the dynamic changes in the brain’s microstructure during this critical period. The atlas includes 60 distinct white matter tracts, including commissural, projection, and association fibers. We employed advanced fetal dMRI processing techniques and tractography to map and characterize the developmental trajectories of these tracts. Our findings reveal that the development of these tracts is characterized by complex patterns of fractional anisotropy (FA) and mean diffusivity (MD), reflecting key neurodevelopmental processes such as axonal growth, involution of the radial-glial scaffolding, and synaptic pruning. This atlas can serve as a useful resource for neuroscience research and clinical practice, improving our understanding of the fetal brain and potentially aiding in the early diagnosis of neurodevelopmental disorders. By detailing the normal progression of white matter tract development, the atlas can be used as a benchmark for identifying deviations that may indicate neurological anomalies or predispositions to disorders.

## 1 Introduction

Fetal life is characterized by a complex and dynamic sequence of neurodevelopmental events. During this period, the brain undergoes exponential growth, neurons reach their cortical and subcortical targets, myelination begins, and transient zones in the telencephalic wall develop and give rise to the mature cerebral hemispheres [1, 2]. These processes form the brain microstructure and give rise to a highly organized connectome [3, 4]. The normal brain development at this stage is also highly vulnerable to various diseases and insults, which can result in lifelong neurodevelopmental and psychiatric disorders [5–7].

Diffusion magnetic resonance imaging (dMRI) is the only non-invasive technique to study brain’s white matter micro-structure and to map its structural connections [8, 9]. The potential of dMRI to investigate these intricate cytoarchitectonic and myeloarchitectonic changes in vivo and to study neurological development, brain function, and brain diseases is enormous [10–15]. Deciphering the functions and dysfunctions of the brain across different stages of life requires a fundamental understanding of its microstructure and structural organization. To achieve this, diffusion metrics such as fractional anisotropy (FA) and mean diffusivity (MD) are widely employed [16]. These metrics are particularly sensitive to changes in the underlying white matter microstructure and are frequently used in studies of brain development [17, 18]. Diffusion MRI can also identify and delineate individual white matter fiber tracts in the brain that enable tract-specific analysis. This is a highly useful capability as micro-structural and macro-structural alterations in specific tracts are commonly used in studying brain development and disorders. Furthermore, building on streamline tractography, dMRI enables mapping of structural connections in the brain. Metrics computed from a graph representation of the structural connectome can be useful for studying brain development and degeneration [19, 20].

Although dMRI has shown tremendous potential in neuroimaging, its application to study the brain in utero has been limited. In particular, despite significant progress in mapping the major white matter fiber bundles using tractography in adults, the developmental trajectories during the fetal stage remain far less explored. This knowledge gap is partly due to the challenges of in-utero imaging, where fetal movement and the inherent limitations of dMRI resolution make it difficult to capture the brain’s rapidly-changing structure. There have been recent strides in fetal dMRI acquisition, preprocessing, and analysis [21–25]. These advances promise to extend the proven postnatal capabilities of dMRI to the intricate and dynamic context of fetal brain development. As a result, an increasing number of studies have employed dMRI to probe the brain micro-structure, study the development of white matter tracts, and assess the structural connectivity in the fetal stage [26–28].

Despite the recent advancements, fetal brain dMRI has remained inferior to its post-natal and adult counterparts in terms of accuracy, reliability, and detail [29–31]. In addition to the image acquisition challenges and low data quality mentioned above, a major source of difficulty in analyzing fetal brain dMRI data is a lack of reference baselines and resources such as atlases. Brain atlases have proven invaluable for studying and characterizing normal and abnormal brain development and for carrying out various challenging neuroimaging analysis tasks. Atlases have been a cornerstone of computational neuroimaging for the past three decades [32, 33]. Given their unique potential, recently, some efforts have been made to create atlases of the fetal brain. A recent survey of existing fetal brain atlases can be found in [34]. Most exiting fetal brain MRI atlases are based on anatomical MRI [35, 36]. A few studies have developed dMRI atlases that map the spatio-temporal changes in diffusion tensor [37] and fiber orientation distribution [38, 39]. These atlases have been used to perform various analyses and extract valuable new insights about fetal neurodevelopment [40–42].

However, there is a lack of a comprehensive *fetal brain white matter tract atlas* to characterize the rapid and complex changes occurring during gestation. To date, such atlases have only been developed for adult brains [43, 44], and some previous neonate studies [45, 46]. Given the vast differences between fetal and pediatric/adult white matter macrostructure and microstructure [47, 48], it is impossible to use an atlas of adult or neonatal brain as a reference for fetal studies. Using an age-mismatched atlas will result in erroneous analysis and limited reproducibility and reliability. An atlas of white matter tracts developed specifically from fetal brain scans will have several crucial benefits, some of which are briefly mentioned below.

- It can portray normal development of white matter tracts in utero. Therefore, it can be used to identify and quantitatively characterize atypical development. By identifying abnormal brain growth patterns at their earliest stage, a normative atlas can facilitate selection of highrisk fetuses for neuroprotective intervention or prenatal clinical counseling. This will lead to accurate personalized information and improved management or treatment of the diseases.
- It can aid in performing a range of important brain image analysis tasks with fetal dMRI data that are currently difficult or impossible to carry out. For example, such an atlas will enable automatic extraction/delineation of specific tracts in individual fetal brains. This is a very common analysis in adult brains, which is commonly performed by comparing a subject’s tractogram with an atlas [49]. This capability will make it possible to isolate and study individual fiber pathways of interest [12, 50].
- A fetal white matter tract atlas can significantly improve automatic analysis of tractography results. This is especially relevant because individual fetus’s whole-brain tractography can generate an enormous number of streamlines with very high false positive rate that are not immediately useful to clinicians or researchers. To eliminate the need for time-intensive manual processing, a white matter atlas can be used to automatically filter and segment streamlines generated with standard tractography algorithms [51–55]. This is especially beneficial when studying large datasets such as in population studies or in clinical settings.
- It will advance our understanding of brain development in utero. It will aid in depicting the brain’s structural connectivity network, provide a reference for brain cortical parcellation, serve as a valuable reference and educational resource [56, 57], and facilitate research and data integration [58]. For instance, such an atlas can be used to integrate multimodal imaging data, combining structural MRI, dMRI, and functional MRI (fMRI) to construct integrated brain atlases that reflect brain morphology, complex fiber architecture, and intrinsic functional organization within a standardized coordinate system [59].
- White matter atlases, either voxel-based binary or probabilistic, have been proposed as a way to determine if a specific tract is present in a particular location in the brain. The advantage of such atlases is that they provide a priori anatomical information that can be utilized for individual subjects. This facilitates the identification of specific tracts without the need for extensive manual annotation [51, 60].

Therefore, the goal of this study was to construct a detailed atlas of white matter tracts in the fetal period. To this end, we leveraged advanced MRI techniques and detailed anatomical knowledge to map the evolving white matter pathways throughout gestation. Additionally, we extended earlier virtual dissection protocols based on adult tractography to address the fact that previous studies on fetal brain have relied on a dissection protocol that has not been explicitly detailed and, hence, are operator-dependent. By filling these gaps, our work aspires to contribute a uniquely valuable resource to neuroscience and clinical practice, enhancing our capacity to understand and address developmental brain disorders from their earliest onset. We complement our work by providing our anatomical descriptions with an open-source group fetal tract atlas in the MNI space [61].

## 2 Materials and Methods

### 2.1 Subjects

This research study utilized data collected from fetal MRI scans conducted in utero. The scans were part of a prospective study conducted at Boston Children’s Hospital (BCH) between March 2013 and May 2019. All participants provided written informed consent, and the BCH Institutional Review Board (IRB) approved the study while following HIPAA guidelines. The inclusion criteria for the study were mothers between 18 and 45 years of age who had normal pregnancies with gestational ages between 22 and 37 weeks. Exclusion criteria included any contraindications to MRI, high-risk pregnancies, fetal central nervous system anomalies, and maternal comorbidities such as diabetes, hypertension, or substance abuse.

### 2.2 Image data acquisition

The MRI scans were conducted using 3.0 T scanners (Skyra and Prisma, Siemens Medical Solutions, Erlangen, Germany) fitted with 16-channel body array and spine coils. The field of view (FOV) and the number of slices captured were determined based on the size of the maternal and fetal subjects. Sedation was not administered to any of the women who were imaged.

Structural fetal brain MRI involved the acquisition of multiple T2-weighted Half-Fourier Single Shot Turbo Spin Echo (HASTE) sequences in each orthogonal plane. Acquisition parameters were: TR: 1400-2000 ms, TE: 100-120 ms, in-plane resolution: 0.9-1.1 mm, and slice thickness: 2 mm with no inter-slice space. For diffusion-weighted images, we obtained 2-8 echo-planar diffusion-weighted image acquisitions, each along orthogonal planes with respect to the fetal head. Each acquisition included one or two b=0 s/mm^2^ images and 12-24 diffusion-sensitized images at b=500 s/mm^2^. Acquisition parameters were: TR: 3000-4000 ms, TE: 60 ms, in-plane resolution: 2 mm, and slice thickness: 2-4 mm.

### 2.3 Data preprocessing

We assessed all imaging studies to ensure they had at least two diffusion-weighted scans free of significant motion-induced artifacts, distortion, and signal loss that could hinder successful motion correction and registration between diffusion-weighted and structural T2-weighted scans. To process each subject’s diffusion and structural MRI data, we used a previously validated pipeline to reconstruct structural and diffusion-weighted volumes with a Kalman filtering-based motion tracking and slice-to-volume registration algorithm [37]. The pipeline registers the dMRI measurements for a subject into a standard atlas space and produces consistent measurements that consist of scattered q-space data in each voxel. For the reconstructed dMRI data, we opted for an isotropic voxel size of 1.2 mm.

We excluded subjects with significant reconstruction errors, insufficient visibility of brain structures, or unusually low data quality. An expert made this decision by visually inspecting mean diffusivity and color fractional anisotropy maps, as well as diffusion tensor glyphs, to ensure the proper direction of the principal eigenvectors.

### 2.4 Estimation of local fiber orientation and tractography

Accurately estimating the local fiber orientation is crucial for tracing neural pathways with streamline tractography. Although advanced techniques are available for analyzing complex fiber structures in adult brains, applying them to fetal brain scans remains challenging due to lower data quality. Hence, simpler tensor models are typically used for fetal dMRI to estimate fiber orientations, as they provide more reliable results with typical fetal dMRI data. In this study, we utilized a weighted linear least squares method to estimate the diffusion tensor in each voxel, followed by computing the diffusion orientation distribution function (dODF) from the diffusion tensor and sharpening the estimated dODF. We used a validated deep learning method to compute an accurate segmentation of the brain tissue into white matter, cortical and subcortical gray matter, and cerebrospinal fluid. The resulting fiber orientation map and tissue segmentation map were used to compute anatomically constrained tractography [62] for each fetus included in this work. Details of the tractography method can be found in [63].

The accuracy of reconstructing different tracts can vary depending on the selected angle threshold. For example, tracts with significant curvatures, such as the optic radiation (OR) [64] or the middle cerebellar peduncle (MCP) may not be accurately reconstructed if the maximum angle threshold is set too low. Conversely, tracts that are mostly straight, such as the frontopontine tract (FPT), may produce inaccurate intra-tract streamline turns if the angle threshold is set too high. To account for this, we generated three separate tractograms for each subject using three different angle thresholds: 15*^◦^*, 20*^◦^*, and 25*^◦^*. Each tractogram consisted of 5 million streamlines.

We aimed to reconstruct various projection, association, and commissural fibers while minimizing the short U-fibers that do not contribute to these major tracts. Since the valid range of streamline lengths varies with brain size, we determined the minimum and maximum streamline lengths based on the whole brain volume (in *mm*^3^) [63]. This approach allowed us to decrease overly long erroneous streamlines while also reducing the presence of short association fibers, which usually do not contribute to major fiber bundles. We used the equations below to calculate the minimum and maximum allowable streamline lengths for each subject.

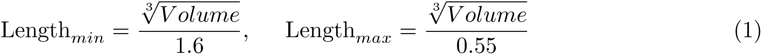

### 2.5 Construction of spatiotemporal diffusion tensor atlas

We used a diffusion tensor-based deformable registration approach to align data from fetuses of the same age. This step is needed for aligning streamline tractography data from different subjects into a common space in order to construct tract atlases. We used the registration methods from the DTI-TK software [65]. For each gestational week between 23 and 36, we considered subjects that were within one week of the target age. For example, all fetuses between 30 and 32 gestational weeks were used to construct an atlas for week 31. Diffusion tensor images *I_i i_*_=1:*N*_ for the *N* fetuses in an age bracket were aligned together using a succession of rigid, affine, and deformable registrations. We denote the combined deformation field to align each *I_i_* to the common space with *ϕ_i_*, as shown in Figure 1.

**Figure 1:**
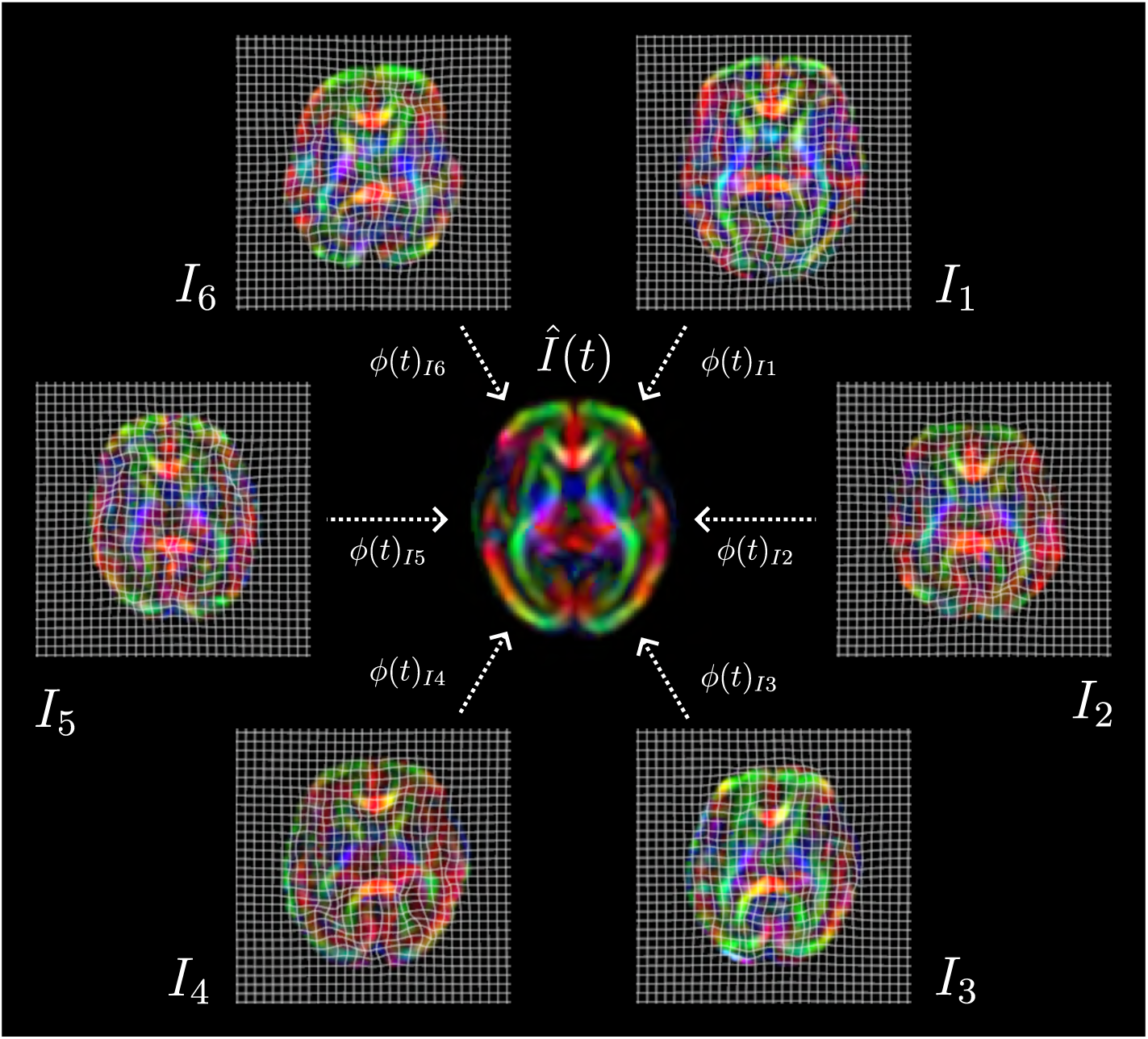
In order to compute spatio-temporal atlases of white matter tracts, we needed accurate alignment of data from multiple fetal brains of the same gestational age. We used deformable registration based on diffusion tensor images *{I_i_}_i_*_=1:*N*_ to compute these deformations, which we denote with *{ϕ_i_}_i_*_=1:*N*_. The schematic in this figure shows an example with six subjects.

### 2.6 Generation of subject tracts

#### 2.6.1 Parcellation of the fetal cortex

In order to extract the white matter tracts of interest from each subject’s tractogram we also required accurate cortical parcellations. For this purpose, we utilized a publicly available parcellated T2-weighted atlas of the fetal brain (http://www.crl.med.harvard.edu/research/fetal_brain_ atlas/). This atlas contains 78 distinct cortical parcellations derived from the Edinburgh Neonatal Atlas (ENA33) [66]. We modified these cortical parcellations by assigning the anterior part of the rolandic operculum to the precentral gyrus and the posterior part to the postcentral gyrus. Additionally, we subdivided the middle frontal gyrus into caudal and rostral portions.

We utilized diffeomorphic image registration [67] to align each age-equivalent T2 atlas to the corresponding MD map of our diffusion tensor atlas. Subsequently, we combined the resulting deformation field with the inverse warp fields obtained during the diffusion tensor atlas computation. Then, the original age-equivalent T2 parcellation was propagated to each subject using a generic interpolator for labeled images [68] (Figure 2).

**Figure 2:**
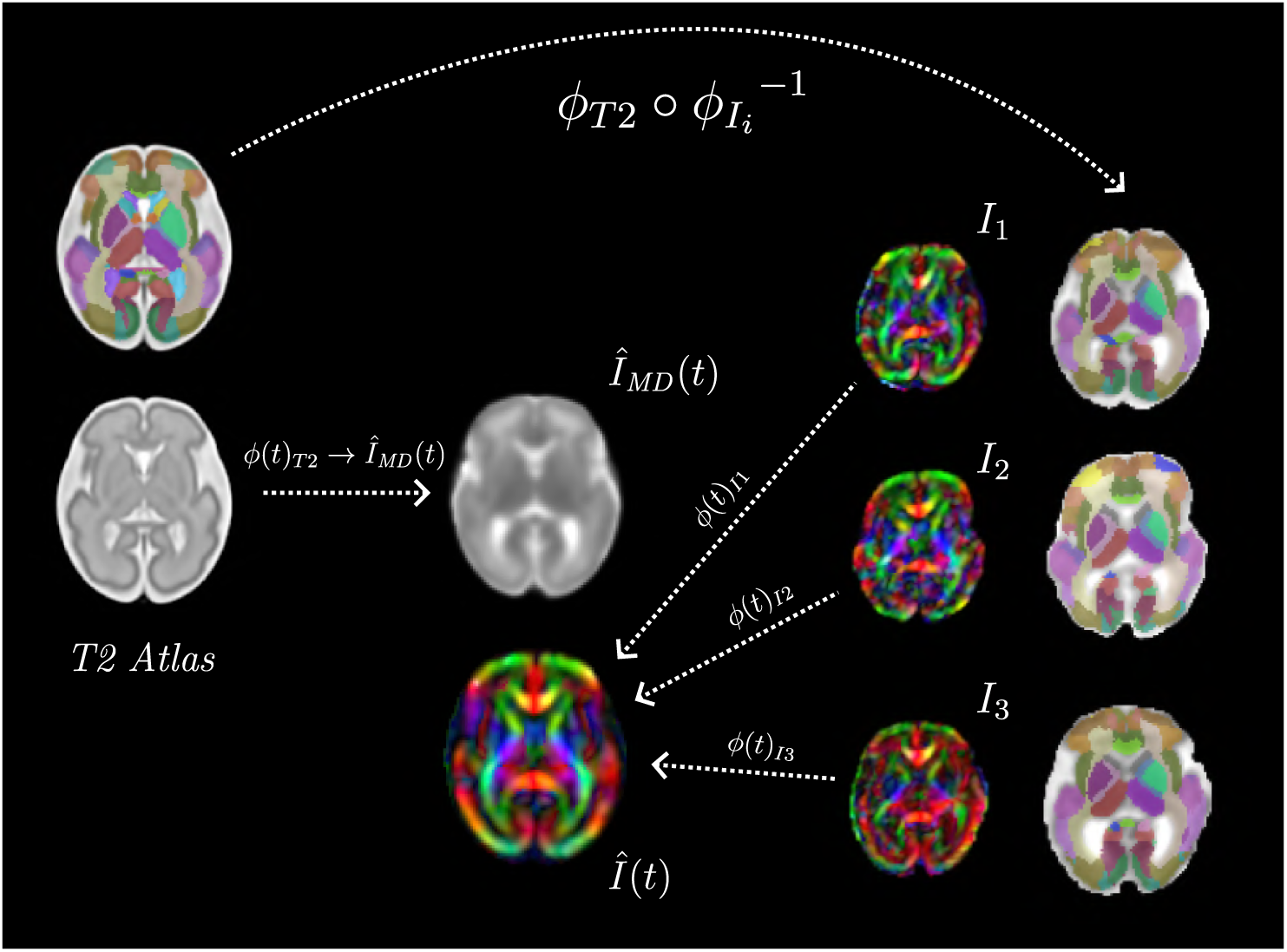
To obtain accurate cortical parcellations for each subject, we registered a T2-weighted spatiotemporal atlas of the fetal brain with cortical parcellation labels to each age-equivalent mean diffusivity map of our generated diffusion tensor atlas. The resulting deformation field was then combined with the inverse warp fields *ϕ^−^*^1^ *_i_*_=1:*N*_ obtained during the diffusion tensor atlas computation to propagate the age-equivalent T2-weighted parcellation to each subject.

#### 2.6.2 Tract extraction

We utilized TractQuerier [69] to extract 60 distinct tracts (see tract descriptions in Section 3.2). TractQuerier extracts tracts based on the streamlines’ starting and ending regions and areas where they should or should not traverse. We defined these regions using the ENA33 parcellations [66], which had previously been modified to suit a T2 spatiotemporal atlas of the fetal brain [35]. We further modified these parcellations as described in Section 2.6.1. We used the queries given by Wassermann et al. [69] and made minor changes to reflect the differences between base parcellation protocols because Wassermann et al. used the FreeSurfer cortical parcellations for adults.

Since we generated three distinct tractographies for each subject (with three different threshold angles), an expert examined each tract and determined the optimal threshold angle for each. Additionally, if a tract from a subject was not reconstructed accurately, it was removed from the analysis. If more than 25% of a subject’s tracts (15 tracts) were found to be inaccurately reconstructed, that subject was excluded entirely from the study.

### 2.7 Construction of the spatiotemporal white matter tract atlas

We used the deformations computed based on diffusion tensor maps, described in Section 2.5, to align the streamline tractography data for each subject into a common space. This was done by applying the deformation fields {*ϕ_i_*}*_i_*_=1:*N*_ to the coordinate points describing each streamline. Let us denote the set of aligned data for a specific tract with *S_i i_*_=1:*N*_, where *S_i_* is the set of tract streamlines for subject *i*. In order to compute an average tract representation, we filtered these streamlines in two steps (Figure 3).

- The first step was intended to remove data from subjects that were not consistent with other subjects in terms of the considered tract. Each subject’s streamline data were converted to a binary mask. This was done by computing the tract density map and removing voxels with tract density below the 5^th^ percentile of non-zero tract density values in order to remove voxels with spurious streamlines. The resulting masks *M_i i_*_=1:*N*_ were merged using the STAPLE algorithm [70] to compute an average tract mask *M̄*. We computed the agreement between each subject’s tract mask and the mean tract mask using the Dice similarity coefficient (DSC) and removed those subjects with DSC(*M_i_, M̄*) *<* DSC_thr_. We used two different values for the threshold DSC_thr_ = 0.80, 0.90. If any subjects were removed, a new average mask *M̄* was computed, and this process was repeated with the remaining subjects until all remaining subjects were consistent.
- The streamlines for the remaining subjects were simply merged into a single bundle of streamlines. Subsequently, this bundle was filtered to remove less reliable streamlines based on Cluster Confidence Index [71]. Any streamline with CCI *<* CCI_thr_ was eliminated. For each tract, we used six different values of CCI_thr_ based on the distribution of CCI for all streamlines in the bundle. Specifically, CCI_thr_ was set to remove 0, 1, 5, 25, 50, and 75 percentile of the CCI values for all streamlines.

**Figure 3:**
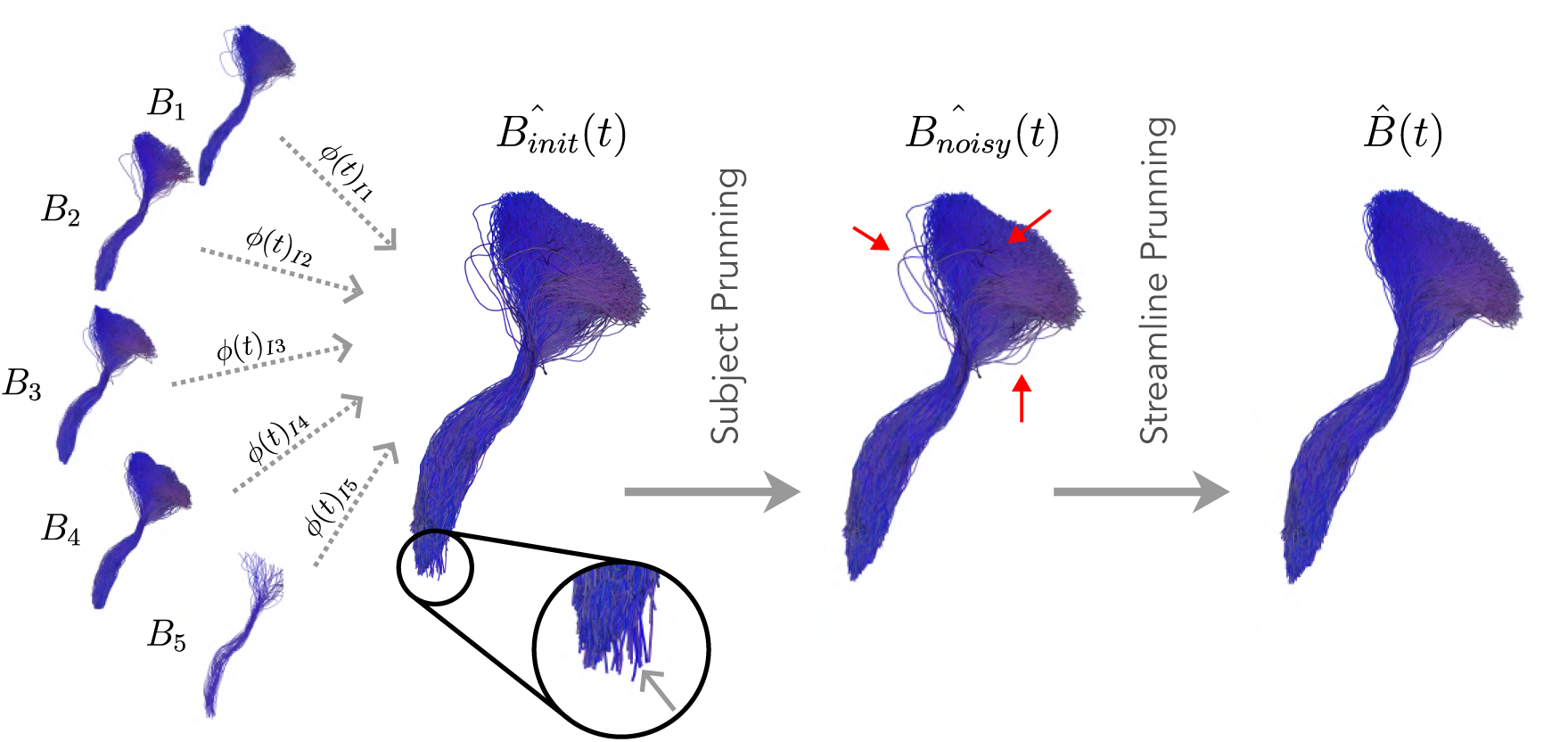
A representative “atlas” tract was computed by merging all the streamlines from multiple subject, separately for each gestational age and tract. The merged tracts were filtered in two steps. First, all streamlines for a subject were removed if the tract mask for that subject was not consistent with an “average mask” computed from all subjects. Second, the streamlines from all remaining subjects were merged into a single bundle of streamlines and the streamlines with low Cluster Confidence Index were eliminated.

With two different values for DSC_thr_ and six different values for CCI_thr_ we obtained 12 different reconstructions for each tract at each gestational age. An expert (C.C., MD, with three years of experience in fetal imaging) visually inspected these 12 reconstructions and selected the one with the fullest coverage and lowest amount of spurious streamlines.

### 2.8 Statistical Analysis

#### 2.8.1 Characterization of the changes in the diffusion metrics with gestational age

For each tract of every subject, we generated a binary mask by computing a density map of streamlines, removing the lowest 1% of non-zero streamline density values, and eliminating any voxels within gray matter structures in order to ensure that our analysis included solely the white matter. Next, we isolated the largest connected component of the binary mask and filled any holes for accuracy. We then calculated median FA and MD values for every tract mask.

We utilized a beta growth and decay model [72] to assess changes in MD over the course of gestation. We opted for this model since earlier studies have identified biphasic changes in MD during development [30, 42]. Furthermore, this model identifies a turning point and can capture the biphasic change that initially rises and then falls with GA. The model is defined by the following equation [72]:

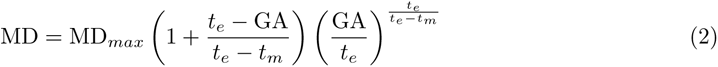

Here, GA represents the gestational age, MD*_max_* is the maximum value of MD attained at time *t_e_*, and *t_m_* is the time of fastest growth rate. To assess changes in FA, based on previous works that have described a biphasic change that first decreases and then decreases with GA [30, 42], we adapted the beta growth and decay model. The modified model is defined as follows:

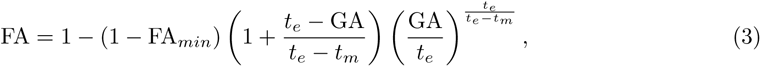

where FA*_min_*is the minimum value of FA, which is reached at time *t_e_*, and *t_m_* is the time of fastest decrease rate.

## 3 Results

### 3.1 Sample characteristics

We included 59 fetal diffusion MRI examinations from 57 distinct fetal subjects. Two of the fetuses were imaged at two different gestational ages. The median gestational age was 29.1 weeks, with a range of 22.6 to 36.9 weeks. Of all the subjects, 35 (58.3%) were male. Figure 4 shows the distribution of gestational age and the number of subjects used to create each atlas.

**Figure 4:**
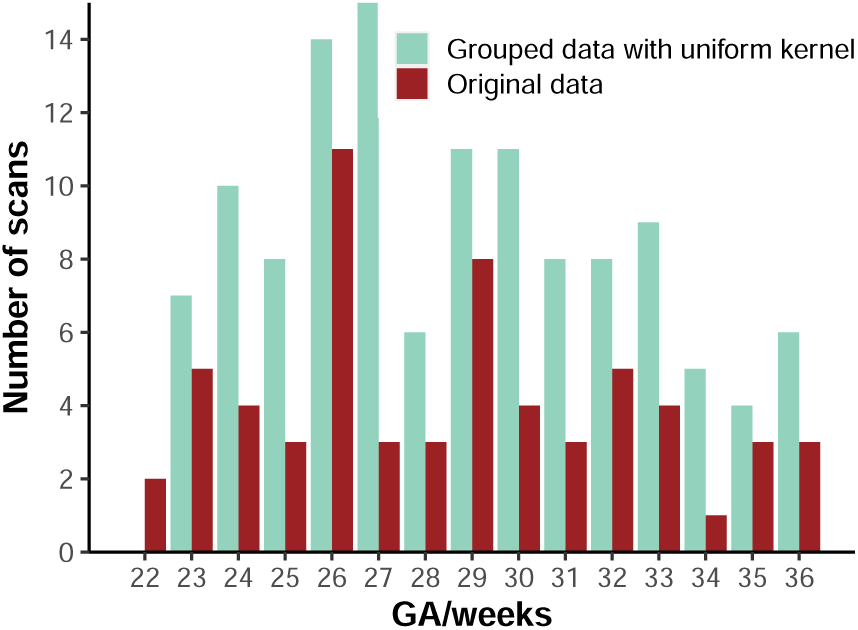
Histogram of the number of fetal scans used at different gestational ages. The red bars show the number of original scans at each GA, while the cyan bars depict the number of subjects used for the atlas construction, as determined using a uniform kernel (GA *±* 1).

### 3.2 Tracts descriptions

A total of 34 tracts were successfully extracted, with 26 having distinct representations in both hemispheres. These tracts were categorized into four classes: association, commissural, projection, and cerebellar. Specifically, we extracted seven commissural tracts, five association tracts, nineteen projection tracts (including nine thalamic, seven striatal, and three “long-range” tracts), and 3 cerebellar tracts. Visual examples of the reconstructed bundles are portrayed in Figure 5. Additional information on the virtual dissection protocol is provided below.

**Figure 5:**
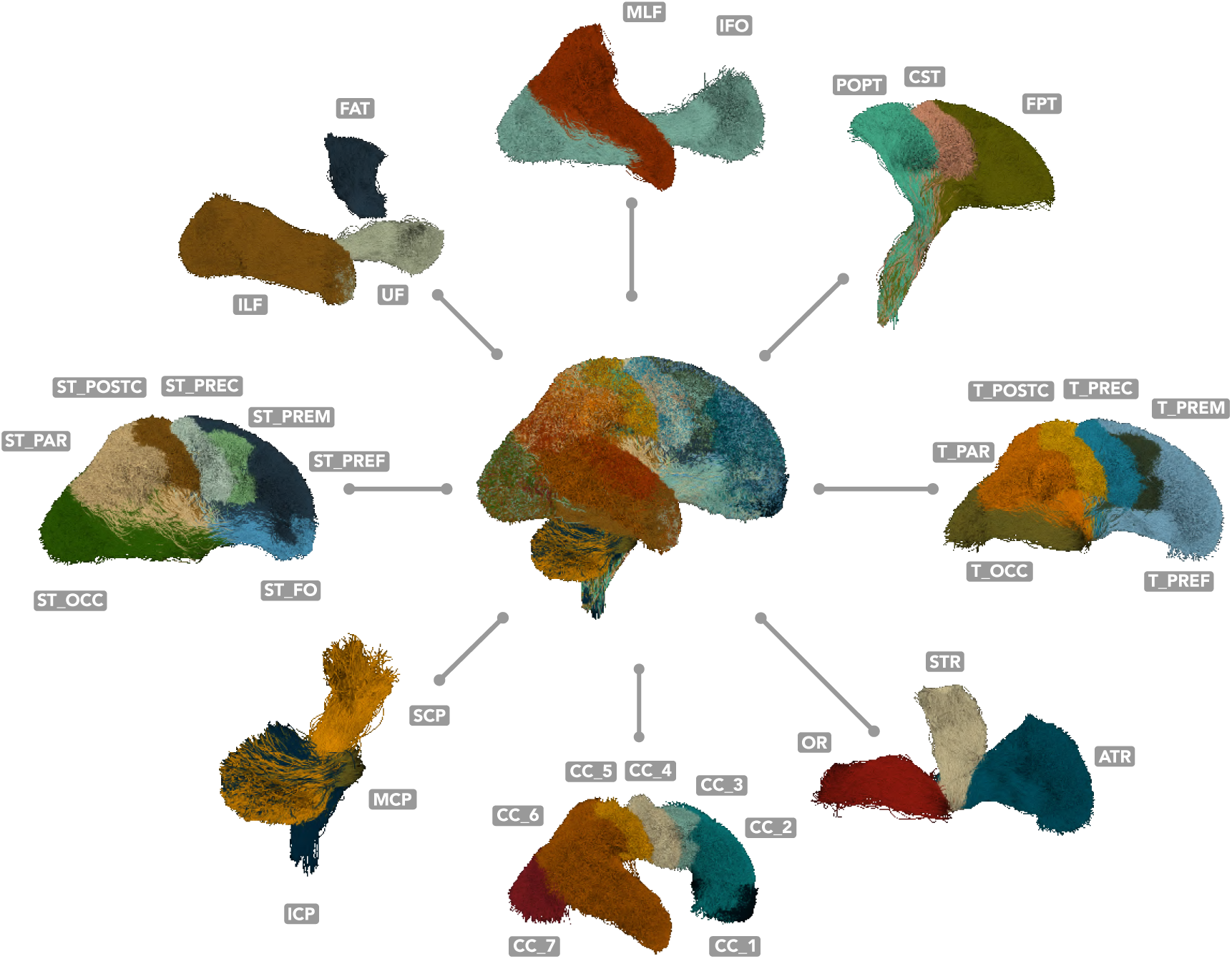
Overview of all generated tracts. For tracts that exist in the left and the right hemispheres, only the right one is shown. The following tracts are included: Anterior thalamic radiation (ATR), Corpus Callosum (Rostrum (CC 1), Genu (CC 2), Rostral body (CC 3), Anterior midbody (CC 4), Posterior midbody (CC 5), Isthmus (CC 6), and Splenium (CC 7)), Corticospinal tract (CST), Frontal Aslant tract (FAT), Fronto-pontine tract (FPT), Inferior cerebellar peduncle (ICP), Inferior fronto-occipital fasciculus (IFO), Inferior longitudinal fascicle (ILF), Middle cerebellar peduncle (MCP), Middle longitudinal fascicle (MLF), Optic radiation (OR), Parieto-occipital pontine tract (POPT), Superior cerebellar peduncle (SCP), Fronto-orbital-striatal (ST FO), Occipito-striatal (ST OCC), Parieto-striatal (ST PAR), Postcentral-striatal (ST POSTC), Precentral-striatal (ST PREC), Prefrontal-striatal (ST PREF), Premotor-striatal (ST PREM), Superior thalamic radiation (STR), Thalamo-parietal (T PAR), Thalamo-postcentral (T POSTC), Thalamo-precentral (T PREC), Thalamo-prefrontal (T PREF), Thalamo-premotor (T PREM), Thalamo-occipital (T OCC), and Uncinate fasciculus (UF).

#### 3.2.1 Commissural fibers

##### Corpus callosum

The corpus callosum (CC) is the largest fiber bundle in the human brain. It facilitates interhemispheric information transfer and integration by connecting the right and left cerebral hemispheres. Traditionally, it has been subdivided into rostrum, genu, body, isthmus, and splenium. In diffusion tensor imaging literature, it is common to find three subdivisions: the genu, forming the forceps minor connecting the left and right prefrontal and anterior cingulate cortices; the callosal body; and the splenium, forming the forceps major connecting the left and right posterior parietal, medial occipital, and medial temporal cortices. Advances in fiber-tracking algorithms and their integration with functional data have led to the development of alternative parcellation strategies. Here, we reconstructed the CC into seven segments by further subdividing the body into the rostral body, anterior midbody, and posterior midbody [73].

##### Rostrum (CC 1)

The rostrum is the anterior-inferior segment of the CC, positioned at the floor of the frontal horn and extending backward to meet the lamina terminalis. It connects the orbital surfaces of the frontal lobes.

##### Genu (CC 2)

The genu forms the anterior bend of the CC. Its fibers curve forward as forceps minor, connecting the medial and lateral surfaces of the frontal lobes. This segment was reconstructed using the bilateral prefrontal cortices and the anterior third of the CC.

##### Rostral body (CC 3)

The majority of the CC comprises the body, where its fibers extend laterally, intersecting with the projection fibers of the corona radiata and establishing connections across wide regions of the cerebral hemispheres. This segment was reconstructed using the bilateral premotor cortices, including the caudal middle frontal gyri, the supplementary motor areas bilaterally, and the central body of the CC.

##### Anterior midbody (CC 4)

This segment was reconstructed using the bilateral primary motor cortices (precentral gyri) and the posterior third of the CC.

##### Posterior midbody (CC 5)

This segment was reconstructed using the bilateral primary sensory cortices (postcentral gyri) and the posterior third of the CC.

##### Isthmus (CC 6)

The isthmus denotes the typical narrowing of the CC at the point where the body meets the splenium. It predominantly carries primary motor, somatosensory, and auditory fibers. This segment was reconstructed using the bilateral parietal cortices, the superior aspect of the bilateral temporal lobes, and the isthmus of the CC.

##### Splenium (CC 7)

The splenium forms the rearmost part of the CC and primarily consists of visual and association temporo-occipital and parietal commissural fibers. The tapetum is formed by fibers of the body and splenium that course around the lateral wall of the ventricular trigones and posterior horns of the lateral ventricles. The fibers interconnecting the occipital lobes curve posteriorly from the splenium, forming the forceps major. The splenium segment was reconstructed using the bilateral occipital cortices and the splenium of the CC [74].

#### 3.2.2 Projection bundles Corticopontine fibers

##### Fronto-pontine tract (FPT)

The fronto-pontine tract (FPT) is part of the corticopontine-cerebellar pathways. It originates in the frontal lobe and projects to the pontine nuclei. These fibers transmit motor signals from the cerebral cortex to the cerebellum, which is essential for coordinating movement. The FPT travels down through the internal capsule (specifically in the anterior half of the posterior limb). These fibers then continue through the cerebral peduncles before synapsing in the pontine nuclei, serving as the relay station where signals are transferred to the opposite cerebellar hemisphere via the middle cerebellar peduncles [75]. This system contributes to the coordination of complex motor functions. Here, we reconstructed the FPT terminating in the brainstem, as ROIs for pontine nuclei were unavailable.

##### Parieto-occipital pontine (POPT)

The parieto-occipital pontine tract (POPT) is part of the corticopontine-cerebellar pathways. It originates in the parietal and occipital lobes and projects to the pontine nuclei. From the pontine nuclei, fibers cross to the opposite side of the brainstem and enter the cerebellum via the middle cerebellar peduncle [76]. The corticopontine-cerebellar pathways are crucial for the integration of sensory information and the coordination of motor functions. Here, we reconstructed the POPT starting from the brainstem, as ROIs for pontine nuclei were unavailable.

##### Corticospinal tract (CST)

The corticospinal tract (CST) is a major pathway that controls voluntary motor function. It originates in the cerebral cortex and passes through the posterior limb of the internal capsule before terminating in the spinal cord. The CST arises from multiple cortical areas, including the primary motor cortex, premotor cortex, supplementary motor area, and somatosensory cortex. Here, we only considered those fibers originating from the primary motor area (precentral gyrus) to avoid overlapping with the FPT and POPT endpoints. Most CST axons cross over at the pyramidal decussation, located at the junction between the brainstem and spinal cord, and continue down the spinal cord in the lateral corticospinal tract. Damage to the CST can result in significant motor deficits.

##### Corticostriatal fibers

Nearly all regions of the cerebral cortex send projections to the striatum, which serves as the gateway to the basal ganglia. Corticostriatal pathways employ various routes to reach their destinations, with many utilizing the external capsule and/or Muratoff’s bundle at some stage. Positioned just lateral to the caudate nucleus, the external capsule curves around the lateral border of the putamen, distinct from its more lateral counterpart, the extreme capsule, which is separated by the claustrum. Muratoff’s bundle lies dorsal to the caudate nucleus, following its upper contour. Studies in monkeys using tract-tracing techniques have revealed fibers destined for the striatum originating from preoccipital cortices, anterior and posterior cingulate cortices, dorsal prefrontal cortex, and other association and limbic areas. While both bundles convey cortical fibers to the striatum, those traversing the external capsule predominantly terminate in the putamen. In contrast, those traversing Muratoff’s bundle tend to terminate in the caudate nucleus. Nonetheless, exceptions to this pattern exist, with axons occasionally crossing between these two bundles [77].

##### Fronto-orbital-striatal (ST FO)

The front-orbital-striatal fibers (ST FO) connect the striatum with the frontal cortex, specifically the orbitofrontal cortex. This tract is part of the larger corticostriatal network, which is implicated in various functions, including decision-making, reward processing, and integrating cognitive and emotional information [78]. Alterations in the topography and connectivity of the ST FO have been linked to various neuropsychiatric illnesses, including obsessive-compulsive disorder [79].

##### Prefrontal-striatal (ST PREF)

The prefrontal-striatal fibers (ST PREF) connect the striatum with the prefrontal cortex. These fibers are part of the corticostriatal pathway, which involves various functions such as motor control, cognitive processes, and emotional regulation. White matter integrity in frontostriatal projections has been linked to impulsivity [80]. At the same time, differences in the structural connectivity of prefrontal-striatal tracts have been shown to mediate age-related differences in action selection [81]. These findings suggest that the integrity of ST PREF fibers plays a critical role in various aspects of behavior and neuropsychiatric conditions.

##### Premotor-striatal (ST PREM)

The premotor-striatal fibers (ST PREM) connect the striatum with the premotor areas of the frontal cortex.

##### Precentral-striatal (ST PREC)

The precentral-striatal fibers (ST PREC) connect the striatum with the primary motor area (precentral gyrus).

##### Postcentral-striatal (ST POSTC)

The postcentral-striatal fibers (ST POSTC) connect the striatum with the primary sensory area (postcentral gyrus).

##### Parieto-striatal (ST PAR)

The parieto-striatal fibers (ST PAR) connect the striatum with the parietal cortex.

##### Occipito-striatal (ST OCC)

The occipito-striatal fibers (ST OCC) connect the striatum with the occipital cortex.

##### Thalamocortical fibers

The thalamocortical radiations are a network of nerve fibers that connect the thalamus to the cerebral cortex. These fibers are critical in transmitting sensory and motor information from the thalamus to specific cortex areas through relay neurons. Each thalamic nucleus is linked to a distinct cortical area through parallel pathways forming thalamocortical radiations. In addition, the corticothalamic fibers relay information from the cortex back to the thalamus. The thalamocortical radiations are essential for transmitting information and help regulate cortical arousal and consciousness. Moreover, through their connections with different cortical regions, they are involved in higher-order cognitive functions [82]. Our work reconstructed thalamic radiations using the whole thalamus as a starting region and the cortical targets as the end regions.

##### Anterior thalamic radiations (ATR)

The anterior thalamic radiation (ATR) connects the dorsomedial, dorsolateral, and anterior thalamic nuclei to the prefrontal cortex, particularly the dorsolateral prefrontal cortex, through the anterior limb of the internal capsule. It is essential in executive functions, planning complex behaviors, memory, and attention. The altered integrity of this tract has been linked to the pathophysiology of neuropsychiatric conditions such as autism spectrum disorders [83] and schizophrenia [84].

##### Thalamo-prefrontal (T PREF)

This bundle was considered an extension of the ATR, but it included endpoints in the entire prefrontal cortex, including the superior frontal gyrus.

##### Thalamo-premotor (T PREM)

The premotor cortex, located just anterior to the primary motor cortex, is involved in planning and organizing movements and actions.

##### Thalamo-precentral (T PREC)

The thalamic motor radiation comprises fiber tracts that either originate or terminate in the ventral lateral nucleus of the thalamus and then extend to the motor cortex. These tracts facilitate communication between the thalamus and the cerebral cortex, particularly regarding motor function. The ventral lateral nucleus is a vital relay station in the motor circuit as it receives input from the cerebellum and transmits output to the motor cortex, playing a crucial role in movement planning and coordination.

##### Superior thalamic radiation (STR)

The superior thalamic radiation (STR) connects the ventral thalamic nuclei to the paracentral lobule through the superior thalamic peduncle, the posterior limb of the internal capsule, and the corona radiata. The STR is crucial in transmitting sensory, motor, and potentially cognitive signals to the cortex.

##### Thalamo-postcentral (T POSTC)

The thalamic-postcentral radiation (T POSTC) originates in the ventral posterior nucleus and relays vital somatosensory information from the body to the primary somatosensory cortex. This information is then processed to provide both tactile perception and proprioception. The integrity of these pathways is essential for proper somatosensory function.

##### Thalamo-parietal (T PAR)

The thalamic-parietal fibers (T PAR) connect the thalamus to the parietal cortex.

##### Optic radiation (OR)

The optic radiation (OR) is a bundle of axonal fibers in the brain that carry visual information from the lateral geniculate nucleus of the thalamus to the primary visual cortex located in the calcarine fissure of the occipital lobe. These fibers are part of the posterior visual pathway and are critical for transmitting visual signals to the cortex. We generated two representations of the optic radiation: one using the classical anatomical definition with endpoints only in the pericalcarine cortex and an extended one that includes endpoints in the entire occipital lobe.

##### Thalamo-occipital (T OCC)

This bundle was considered to be the same as the extended optic radiation, where endpoints are located in the entire occipital lobe, including its lateral aspect.

#### 3.2.3 Association fibers

##### Uncinate fasciculus (UF)

The uncinate fasciculus (UF) connects the anterior temporal lobe with the orbitofrontal cortex. It plays a role in various functions, such as language, episodic memory, and social-emotional processing. The UF has also been linked to the ability to decode facial emotion expressions, suggesting its importance in social behavior [85]. The UF is one of the last tracts to mature, continuing its development into the third decade of life, and individual differences in its maturation may explain behavioral variations [86]. Abnormalities in the UF have been observed in conditions like Phelan-McDermid syndrome, where changes in its microstructure may contribute to deficits in social and emotional interaction [87]. In neurodegenerative diseases, the integrity of the right UF has been linked to socioemotional sensitivity, with damage to this tract affecting socioemotional functioning [88].

##### Frontal Aslant Tract (FAT)

The frontal aslant tract (FAT) connects the pars opercularis and pars triangularis of the inferior frontal gyrus (IFG) to the supplementary motor area (SMA). Extensive research has identified this tract as crucial in multiple facets of speech and language function [89], encompassing verbal fluency, speech initiation and inhibition, and sentence production [90]. Moreover, the FAT is also implicated in executive functions such as working memory, attention, motor control, and social communication [91].

##### Inferior fronto-occipital fascicle (IFOF)

The inferior frontal-occipital fascicle (IFOF) is a prominent association fiber bundle that connects the occipital region with the inferior frontal gyrus, frontal-orbital region, and frontal pole. At the level of the extreme capsule, the IFOF narrows. Although theoretically distinct from other temporal lobe association pathways, the IFOF runs parallel to the middle longitudinal fasciculus, inferior longitudinal fasciculus, and uncinate fasciculus. The IFOF plays a crucial role in various cognitive functions and has been linked to semantic language processing [92], goal-oriented behavior, and facial recognition functions [93].

##### Middle longitudinal fascicle (MLF)

The Middle Longitudinal Fascicle (MLF) is a significant associative pathway that connects the superior temporal gyrus to the parietal and occipital lobes. Its involvement in high-order functions such as language, attention, and integrative higher-level visual and auditory processing, particularly related to acoustic information, has been hypothesized [94]. Additionally, the MLF’s clinical relevance has been identified in several neurodegenerative disorders and psychiatric conditions, including Primary Progressive Aphasia [95] and Schizophrenia [96].

##### Inferior longitudinal fascicle (ILF)

The inferior longitudinal fasciculus (ILF) is a prominent associative tract that connects the occipital and temporal-occipital areas to the anterior temporal regions and plays an important functional role in visual memory and emotional processing. The ILF is involved in various cognitive and affective processes that operate on the visual modality, such as object and face recognition, visual emotion recognition, language, and semantic processing (including reading and memory) [97, 98]. The left ILF has also been linked to orthographic processing, and its disruption may contribute to difficulties in this domain [99].

#### 3.2.4 Cerebellar bundles

##### Inferior cerebellar peduncle (ICP)

The Inferior cerebellar peduncle (ICP) primarily comprises sensory fibers conveying information from the spinal cord to the cerebellum. Functionally, the ICP plays a crucial role in maintaining balance and posture by integrating proprioceptive sensory and motor functions. Microdissection studies have identified four afferent bundles and one efferent bundle within the ICP. However, in our reconstruction, we simplify it into a single lateralized bundle incorporating the cerebellum’s dentate, fastigial, and interposed nuclei, along with the ipsilateral medulla oblongata [73].

##### Middle cerebellar peduncle (MCP)

The Middle cerebellar peduncles (MCPs) are substantial paired bundles linking the brainstem to the cerebellum bilaterally. In neuroimaging studies, they are often depicted as a unified commissural bundle connecting both cerebellar hemispheres. The MCPs are believed to influence the modulation of skilled manual motor functions and have been anatomically delineated into three sub-fascicles (superior, inferior, and deep) via microdissection studies. In our reconstruction, we represented the MCPs for each side, utilizing the contralateral pons and cerebellar cortex. As a result, the streamlines traverse the midline in the pons, aligning with its neuroanatomical definition [73].

##### Superior cerebellar peduncle (SCP)

The superior cerebellar peduncles are paired bundles of white matter fibers that establish connections between the cerebellum and the midbrain. These peduncles are integral conduits for crucial afferent and efferent fibers, comprising pathways such as the cerebellothalamic, cerebellorubral, and ventrospinocerebellar tracts [73].

### 3.3 Changes in white matter diffusivity across gestation

#### 3.3.1 Changes in fractional anisotropy (FA)

As shown in Figure 6, for all tracts, fractional anisotropy followed a biphasic change that first decreased and then increased with GA. The *T_e_* time varied among tracts, with the majority reaching it at some point between 28 and 32 weeks. The CST and POPT were the earliest tracts to reach *T_e_*, while the ATR was the last. The T PREC was the first thalamic tract to reach *T_e_*, while the ST PREC was the first of the striatal tracts. In the corpus callosum, the midbody was the first section to reach *T_e_*, and the rostrum was the last. On average, commissural tracts reached *T_e_* at 30.81 weeks, long projection tracts at 29.25 weeks, thalamic tracts at 30.13 weeks, striatal tracts at 30.49 weeks, and association tracts at 30.56 weeks. The point of lowest FA showed minor variation between tracts, averaging at 0.11. The FATs had the lowest FA values, while the CSTs and POPTs had the highest minimum values among all tracts.

**Figure 6:**
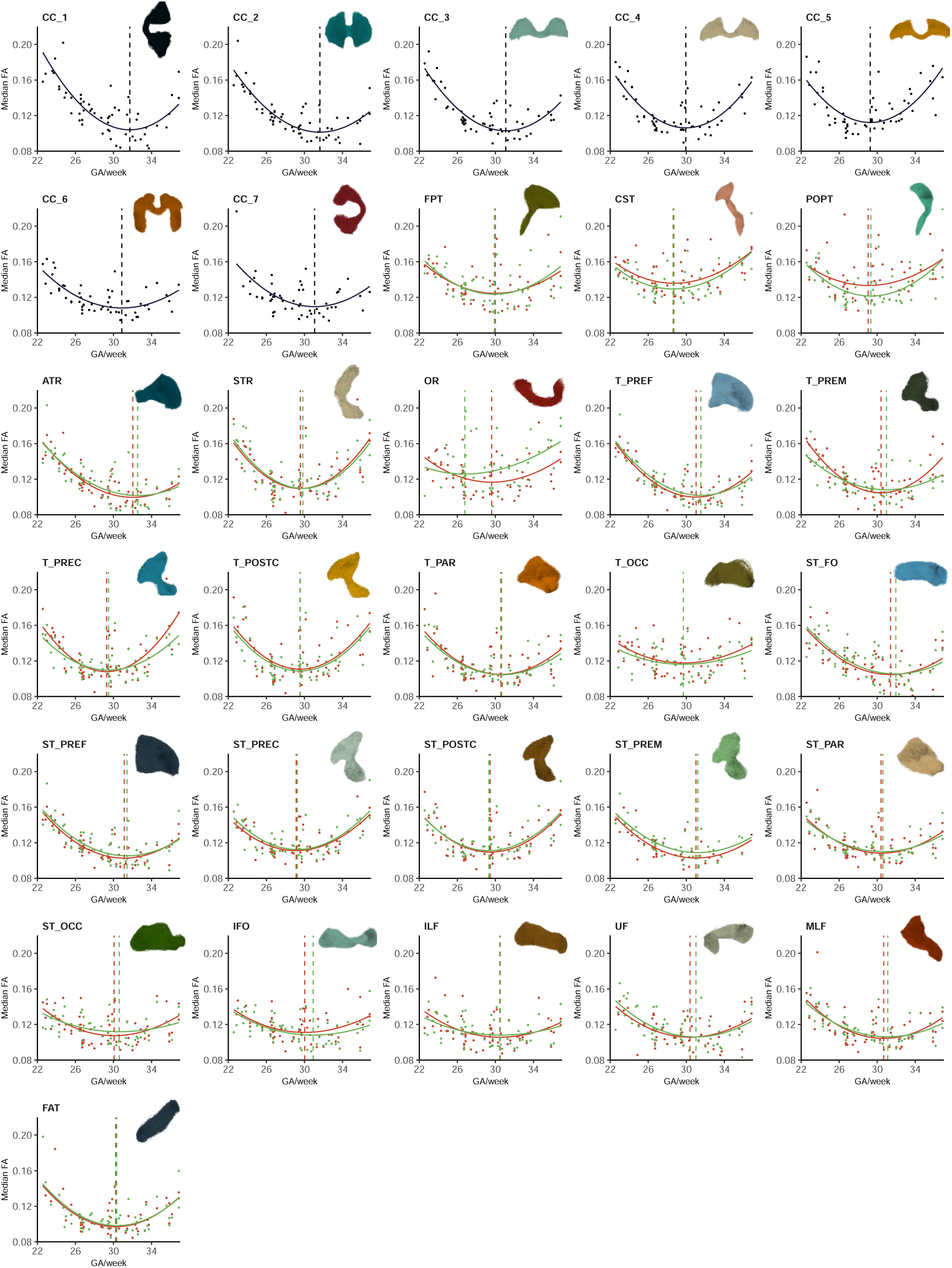
Developmental trends of Fractional Anisotropy (FA) for different tracts of the fetal brain portrayed with a beta growth and decay model. The vertical dashed line indicates *t_e_* in different tracts (unilateral [dark blue], left [red], and right [green] sides). Please refer to the caption of Fig. 5 for the full tract names of abbreviations used in this figure.

#### 3.3.2 Changes in mean diffusivity (MD)

For all tracts, the mean diffusivity followed a biphasic change that initially increased and then decreased with GA as shown in Figure 7. The *T_e_* time varied among tracts, with the majority reaching it at some point between 27 and 30.5 weeks. The CST and POPT were the earliest tracts to reach *T_e_*, while the ST FO was the last. The T POSTC was the first thalamic fiber to reach *T_e_*, while the ST POSTC was the first of the striatal fibers. In the corpus callosum, the midbody was the first section to reach *T_e_*, and the rostrum was the last. On average, commissural tracts reached *T_e_* at 28.88 weeks, long projection fibers at 25.73 weeks, thalamic fibers at 28.52 weeks, striatal fibers at 28.30 weeks, and association fibers at 29.11 weeks. The peak MD exhibited minor variation among tracts, averaging at 1.55 10*^−^*^3^ mm^2^*/*s. The CSTs and POPTs had the lowest MD peaks, while the FATs had the highest among all tracts.

**Figure 7:**
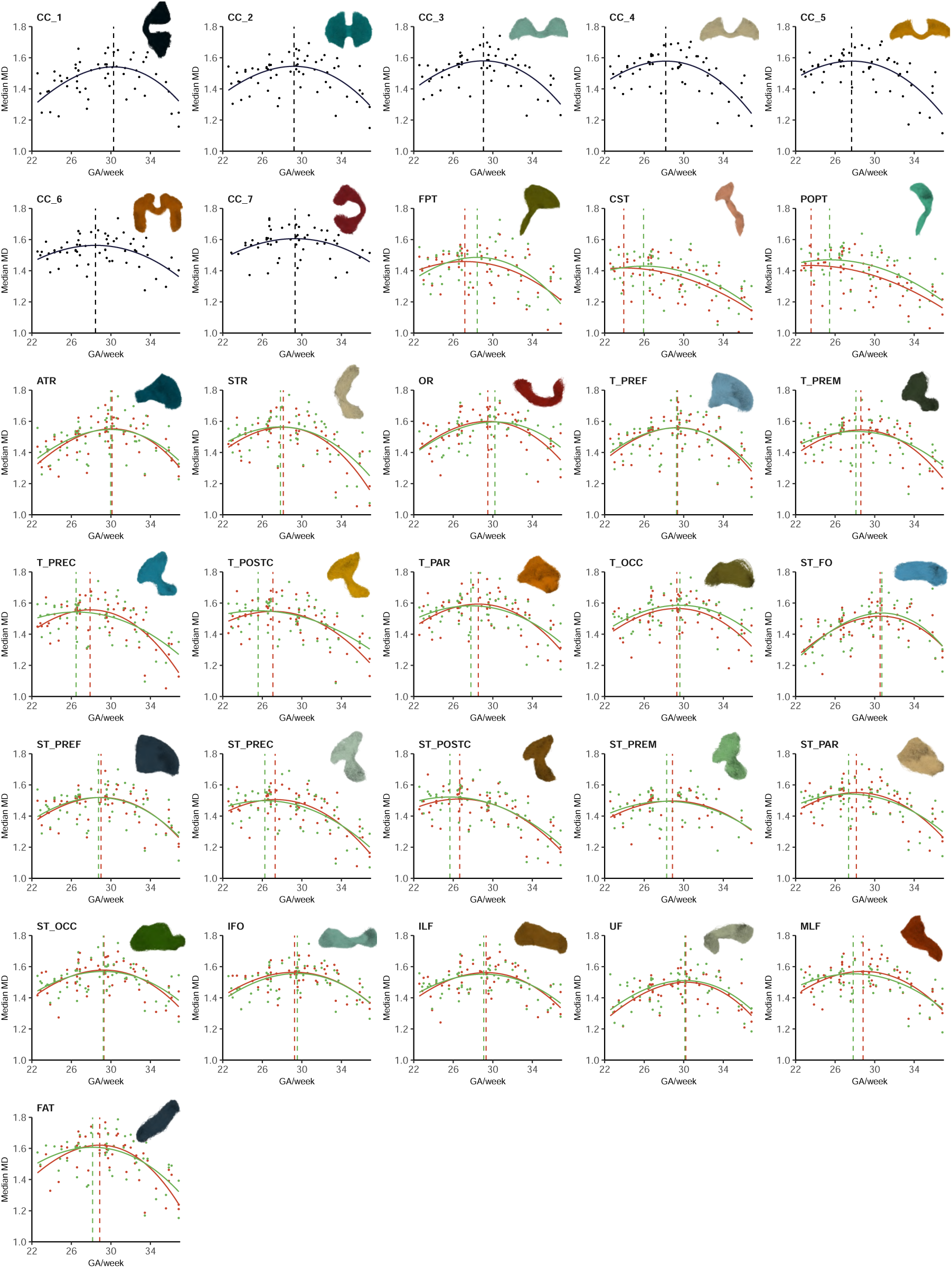
Developmental trends of Mean Diffusivity (MD) (expressed in 10*^−^*^3^ mm^2^*/*s) for different tracts of the fetal brain portrayed with a modified beta growth and decay model. The dashed line indicates *t_e_* in different tracts (unilateral [dark blue], left [red], and right [green] sides). Please refer to the caption of Fig. 5 for the full tract names of abbreviations used in this figure.

## 4 Discussion

In this study, we have meticulously created a detailed atlas of white matter tracts for the fetal brain, covering the span from 23 to 36 weeks of gestational age. Utilizing advanced diffusion MRI, reconstruction, and tractography techniques, we were able to reconstruct 60 different tracts, including commissural, association, and projection tracts. This atlas represents a significant stride in understanding the intricate architecture of the developing brain, offering a glimpse into the dynamic changes occurring in utero.

Our findings also reveal that the developmental trajectories of FA and MD across different white matter tracts exhibit a biphasic pattern in the period studied in this work. These trends reflect the complex interplay of neurodevelopmental processes such as axonal growth, myelination, and synaptic pruning. The variation in the time points at which tracts reach their peak or nadir of FA and MD underscores the heterogeneity in the maturation rates of different brain regions.

Our findings on developmental trends align with previous tractography studies that have reported white matter microstructural metrics derived from the widely used diffusion tensor model. Our study correlates well with the findings of Machado et al. [42] and Wilson et al. [30] for almost all studied tracts, including the forceps minor (CC 1 in our work), CC 2, IFO, ILF, OR and UF showing a biphasic trend in both FA and MD. Those studies found that a second-degree polynomial best-described the changes in FA and MD. The only differences with our study were observed in the splenium in both FA and MD, in the CST in MD, and in the forceps major (CC 7 in our work) in both FA and MD, which exhibited linear trends in those works. On the other hand, previous studies have used study populations with very few subjects under 28 weeks. Thus, the reported linear increase in FA and decrease in MD in the third trimester in those previous studies [40, 100] corresponds to the changes in FA and MD that we observe in our data between approximately 29 weeks and the end of our study period.

Our study’s results closely match the timing and location of significant histologic milestones that take place in late gestation [1]. Due to the sensitivity of tensor metrics to various biological processes, the observed trends are likely influenced by multiple factors. For most tracts, we observed a U-shaped function for FA and an inverted U-shaped fit for MD. The change in trajectory for both functions occurred around the 30th week of gestational age, which coincides with the involution of the radialglial scaffolding that facilitates neuronal migration and precedes the dissolution of the subplate, as seen histologically [101]. Radial glia is known to hinder water diffusion. Thus, its involution leads to low anisotropy and increased diffusivity. Later in gestation, when neuronal migration halts, the scaffold involutes, and the inflow of thalamic axons increases the space between neurons, elevating diffusivity and facilitating a more uniform diffusion of water [101]. These findings are consistent with previous histological studies that have reported a significant decrease in the abundance of extracellular matrix (ECM) between white-matter fiber bundles by 35 weeks of gestation [102]. Given the high water content of ECM, its relative abundance around fibers may contribute to changes in MD values and significantly influence the observed trends.

Although FA is often used as a measure of integrity or myelination in older individuals, the white-matter pathways in this study are still developing at birth. Only after birth are mature myelin markers expressed in the cerebral cortex, with substantial increases occurring in the first two years of life [103]. Despite this, unmyelinated white-matter tracts still exhibit changes in signal intensity consistent with anisotropic water diffusion [104]. Therefore, the observed increases in FA may be the result of active axonal growth and initial ensheathment of axons by premyelin sheaths generated by immature oligodendrocytes [105, 106].

The comprehensive mapping offered by the new atlas developed in this study can enable significant new applications. It enhances our understanding of normal fetal brain development and lays the groundwork for identifying deviations that may signal neurodevelopmental disorders. While previous works have been limited to a small subset of the major fiber pathways, we have successfully reconstructed many of the most representative tracts in the fetal brain. This advancement aligns with postmortem studies suggesting that commissural, association, and projection fibers are present as early as the first trimester of gestation [26]. Nonetheless, we encountered challenges in reconstructing the OR; we could not reliably reconstruct the ICP and SCP, and we could not reconstruct the superior longitudinal fasciculus (SLF) and the arcuate fasciculus (AF).

In our work, the OR had few streamlines reaching the occipital cortical targets in fetuses below 27 weeks of gestation. According to electron microscopy studies, the OR can be observed emerging from the lateral geniculate nucleus to the subplate between 11 and 13 weeks of gestation [107, 108]. However, the presence of synapses in the cortical plate is only observable at 23 to 25 weeks of gestation [108]. Therefore, the challenges we have faced in reconstructing this tract in the younger ages are consistent with the fact that synapses in the cortical plate develop much later in fetal development than in other brain areas.

We also encountered difficulties in accurately reconstructing the SCP and ICP. Our reconstructions generated an SCP in which a significant portion of the streamlines traveled through the middle cerebellar peduncle, and also occupied the whole ipsilateral cerebral peduncle on its way up. Similarly, our reconstruction of the ICP used almost the entire middle cerebellar peduncle and the entire brainstem on its way down. A study by Re et al. [109], where high angular resolution diffusion imaging (HARDI) data on in-vivo, post-natal subjects were used, found that from three subjects, the ICP was sparsely detected in two of the subjects and undetected in the third subject who was a term subject at 40 weeks of gestation. In most subjects aged 2-6 months, the ICP pathways appeared rather sparse and fragmented. These pathways seemed to become more robust in subjects older than six months and branched more extensively in subjects six years and older. Thus, it seems that reconstructing this tract at this early stage is not feasible with the current technology, even more so when using in-utero imaging. Although we provide representations of these tracts in the publicly available atlas, we did not provide developmental trends for these tracts due to being unable to reconstruct them reliably.

Previous studies indicate that SLF tracking may not be feasible, and it may only be partially detectable during the third trimester and later in the neonatal period [26]. The emergence of SLF might occur late during fetal development [26, 28, 110]. Because SLF is one of the more advanced association fibers, it takes longer to mature than other fiber tracts. This means that low levels of myelination during the fetal and neonatal periods [111] may have an impact on the tractography results. Additionally, the SLF intersects with several fiber tracts in its running regions, including the corpus callosum and corticospinal tracts, which can make it more difficult to reconstruct with tractography.

Lastly, we were unable to reconstruct the arcuate fasciculus (AF). According to Horgos et al., [112], the AF is a complex bundle of nerve fibers that is a direct component of the SLF complex. Di Carlo et al. reported that their dissection of the AF/SLF complex was limited in their specimens, leading to the idea that these connections are largely developed postnatally, even from an anatomical perspective [113]. This development timeline suggests that the AF may not be fully formed or too immature to be detected using current tractography techniques during earlier gestational ages, as reported by Huang et al. [26].

Despite the advancements presented in our study, several limitations warrant mention. First, the resolution and signal-to-noise ratio of in-utero diffusion MRI remain inferior to postnatal imaging capabilities, which can restrict the fine detail. The implications of this limitation are most evident in our inability to detect or accurately reconstruct certain tracts, particularly in younger gestational ages. Second, the current analysis relies heavily on the assumptions and simplifications of diffusion tensor imaging, which may not fully capture the complex fiber configurations in certain regions of the developing brain, where white matter tracts cross or diverge. Furthermore, our methodological approach, particularly the streamline tractography, might introduce biases such as false positives or false negatives in tract reconstruction, which can affect the anatomical accuracy of our atlases. Finally, our sample, though larger than those in many previous studies, still represents a relatively small cohort, predominantly composed of subjects from a single population (in our case, only from Massachusetts, USA), which may not provide a complete picture of developmental variability.

## 5 Conclusion

In conclusion, our work’s clinical implications are profound. The atlas serves as a valuable reference for clinicians, researchers, and educators, aiding in interpreting fetal neuroimaging studies and fostering early detection and intervention strategies for brain anomalies. Moreover, this atlas can facilitate the integration of fetal neuroimaging data across studies, enhancing our ability to understand and characterize the impact of environmental, genetic, and maternal health factors on brain development.

## Acknowledgements

This research was supported in part by the National Institute of Neurological Disorders and Stroke under award numbers R01NS128281 and R01 NS106030; the Eunice Kennedy Shriver National Institute of Child Health and Human Development under award numbers R01HD110772 and R01 HD109395; the National Institute of Biomedical Imaging and Bioengineering under award numbers R01 EB031849, R01 EB032366, R01 EB018988, and R01 EB013248; the Office of the Director of the NIH under award number S10 OD025111; the National Science Foundation (NSF) under grant number 212306; the Rosamund Stone Zander Translational Neuroscience Center, Boston Children’s Hospital; the Office of Faculty Development at Boston Children’s Hospital; and a scholarship from the American Roentgen Ray Society. This research was also partly supported by NVIDIA Corporation and utilized NVIDIA RTX A6000 and RTX A5000 GPUs. The content of this publication is solely the responsibility of the authors and does not necessarily represent the official views of the NIH, NSF, or NVIDIA.

